# Six-state amino acid recoding is not an effective strategy to offset the effects of compositional heterogeneity and saturation in phylogenetic analyses

**DOI:** 10.1101/729103

**Authors:** Alexandra M. Hernandez, Joseph F. Ryan

**Author notes:** Corresponding author: Joseph F. Ryan.

## Abstract

Six-state amino acid recoding strategies are commonly applied to combat the effects of compositional heterogeneity and substitution saturation in phylogenetic analyses. While these methods have been endorsed from a theoretical perspective, their performance has never been extensively tested. Here, we test the effectiveness of 6-state recoding approaches by comparing the performance of analyses on recoded and non-recoded datasets that have been simulated under gradients of compositional heterogeneity or saturation. In all of our simulation analyses, non-recoding approaches greatly outperformed 6-state recoding approaches. Our results suggest that 6-state recoding strategies are not effective in the face of high saturation. Further, while recoding strategies do buffer the effects of compositional heterogeneity, the loss of information that accompanies 6-state recoding outweighs its benefits, even in the most compositionally heterogeneous datasets. In addition, we evaluate recoding schemes with 9, 12, 15, and 18 states and show that these all outperform 6-state recoding. Our results have important implications for the more than 70 published papers that have incorporated 6-state recoding, many of which have significant bearing on relationships across the tree of life.

## Introduction

Compositional heterogeneity and substitution saturation are major challenges to phylogenetic inference. Compositional heterogeneity stems from the tendency of genes or organisms to have unequal proportions of amino acids (Collins et al. 1994; Foster and Hickey 1999). These unequal amino acid frequencies are caused by mutational and selective pressures acting at the nucleotide level (Singer and Hickey 2000; Knight et al. 2001), as well as differences in translational efficiency (Akashi and Eyre-Walker 1998). The combination of evolutionary and biological processes results in different amino acid compositions across taxa on the tree. Consequently, challenges to phylogenetic analyses arise when distantly related taxa share sequence similarities due to homoplasy (convergence), rather than descent from a common ancestor (Foster and Hickey 1999; Tarrío et al. 2001).

Similarly, phylogenetic reconstruction artifacts emerge under substitution saturation of amino acids. Substitution saturation occurs when there have been multiple amino acid substitutions at the same site washing out the evolutionary signal (Ho and Jermiin 2004). Like compositional heterogeneity, sequence saturation can lead to long branch attraction, driving unrelated taxa to group together in a clade due to homoplasy (Felsenstein 1978; Hendy and Penny 1989).

Matrix recoding has been proposed as a solution for both compositional heterogeneity and substitution saturation. Under matrix recoding methods, nucleotides or amino acids are recoded into groups based on function (Blanquart and Lartillot 2006). For example, under the RY nucleotide recoding strategy, purines (i.e., A and G) are coded with the character R and pyrimidines (i.e., T and C) are coded with the character Y (Woese et al. 1991; Phillips et al. 2001). In this recoding scenario, only transversion events are meaningful in a phylogenetic analysis. A similar recoding strategy has been implemented for amino acids, the most well-known being Dayhoff 6-state recoding. In Dayhoff 6-state recoding, chemically related amino acids that frequently replace each other are pooled together into six groups based on similar substitution scores in the Dayhoff (or PAM250) matrix (Dayhoff et al. 1978): AGPST, DENQ, HKR, ILMV, FWY, and C (Embley et al. 2003; Hrdy et al. 2004). Thus, only amino acid changes between categories, and not within categories, are considered substitutions. Since the introduction of Dayhoff 6-state recoding, several other 6-state amino acid recoding strategies based around other scoring matrices have been developed. For example, S&R 6-state recoding (Susko and Roger 2007; Feuda et al. 2017) is based on the JTT matrix (Jones et al. 1992) and KGB 6-state recoding (Kosiol et al. 2004; Feuda et al. 2017) is based on the WAG matrix (Whelan and Goldman 2001).

To date, there are at least 77 phylogenetic studies that have implemented six-state amino acid recoding strategies (Table 1). While amino acid recoding has been valued from a theoretical perspective, the performance of 6-state recoding has never been tested empirically against non-recoding methods. In this study, we simulate datasets with either a gradient of compositional heterogeneity or saturation and compare the performance of maximum-likelihood analyses on 6-state recoded datasets to the same analyses on non-recoded datasets. We also run a subset of these analyses using 9-, 12-, 15-, and 18-state recoding schemes and compare these results to those achieved with 6-state recoded and non-recoded matrices.

**Table 1.**
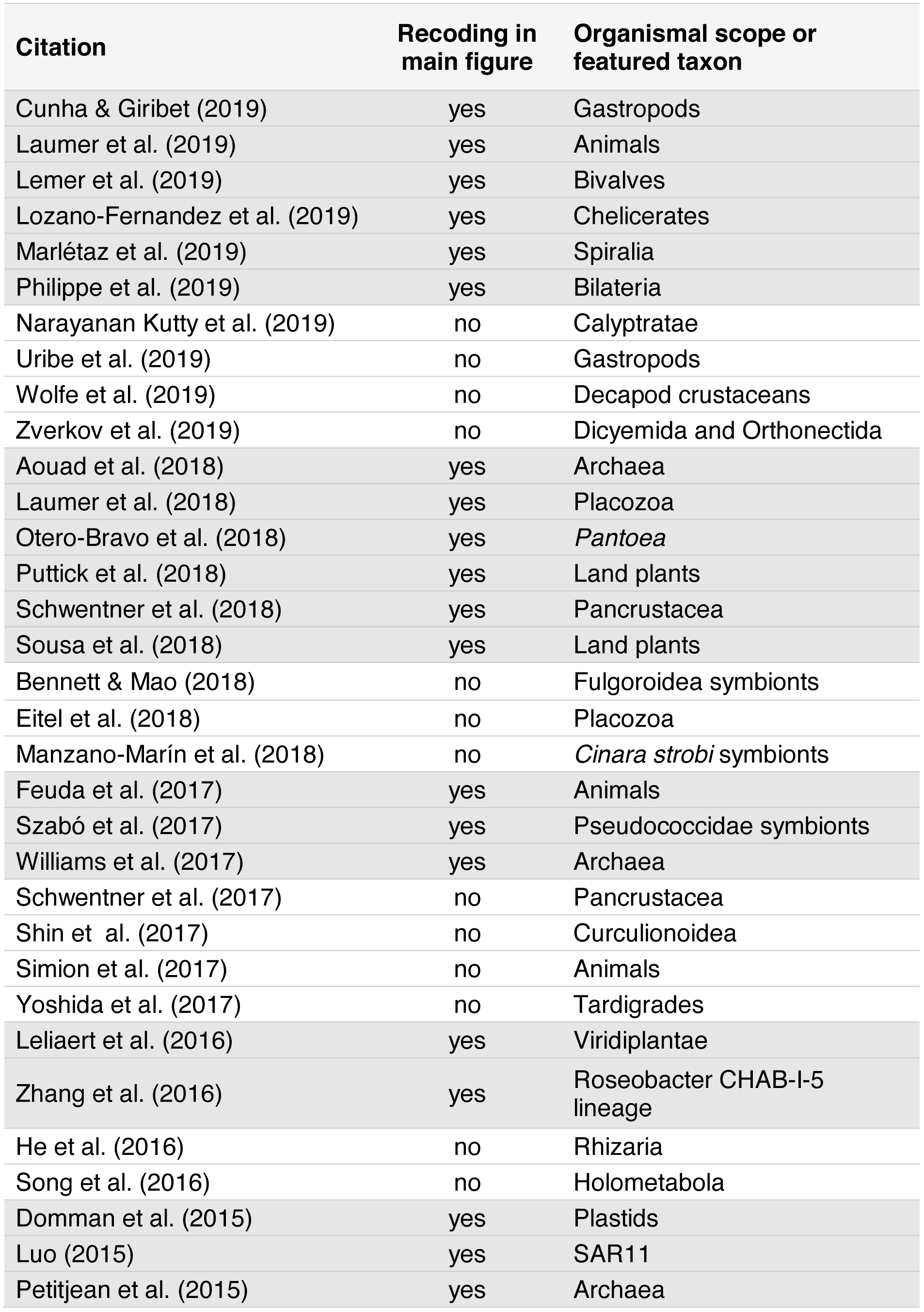
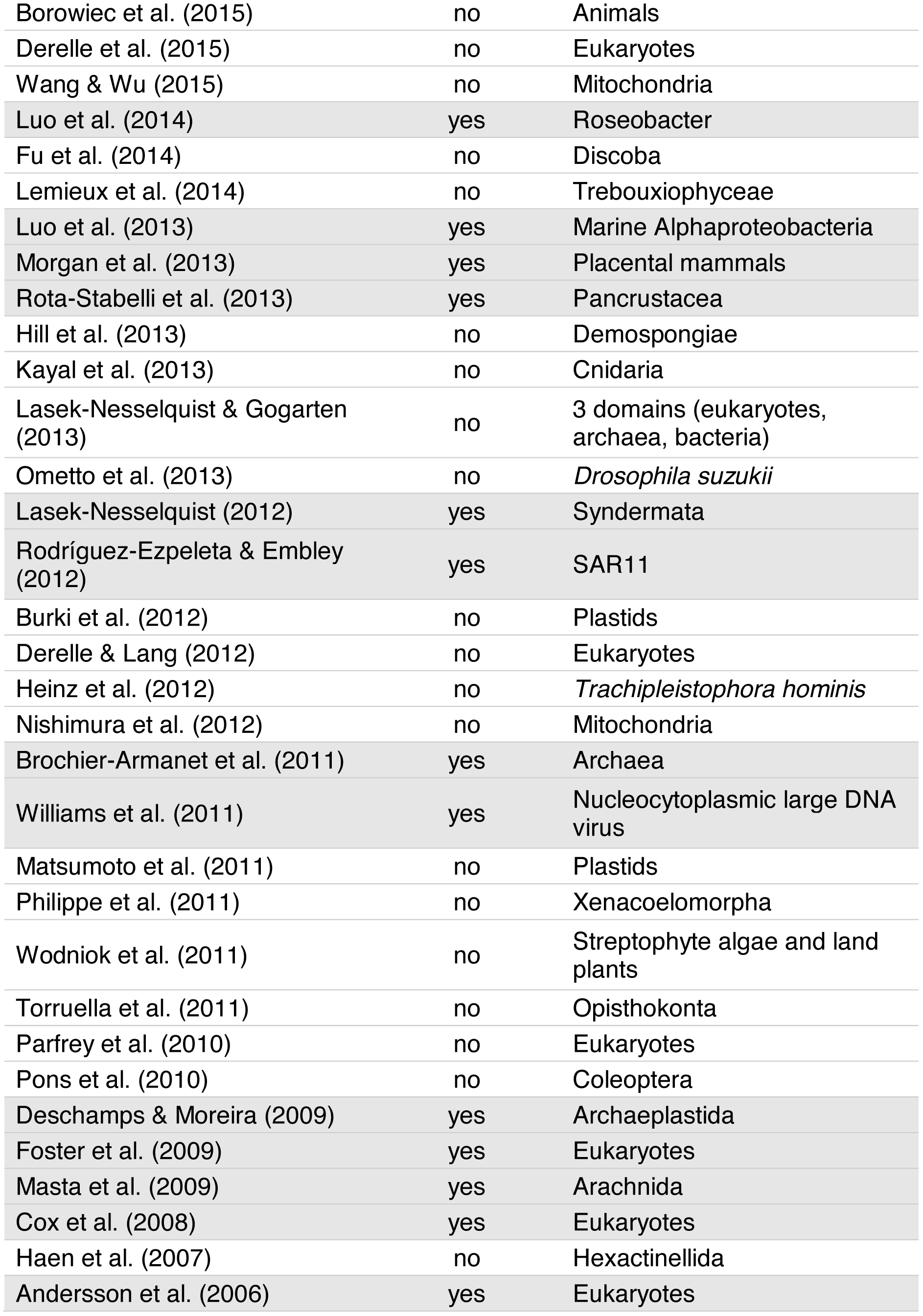
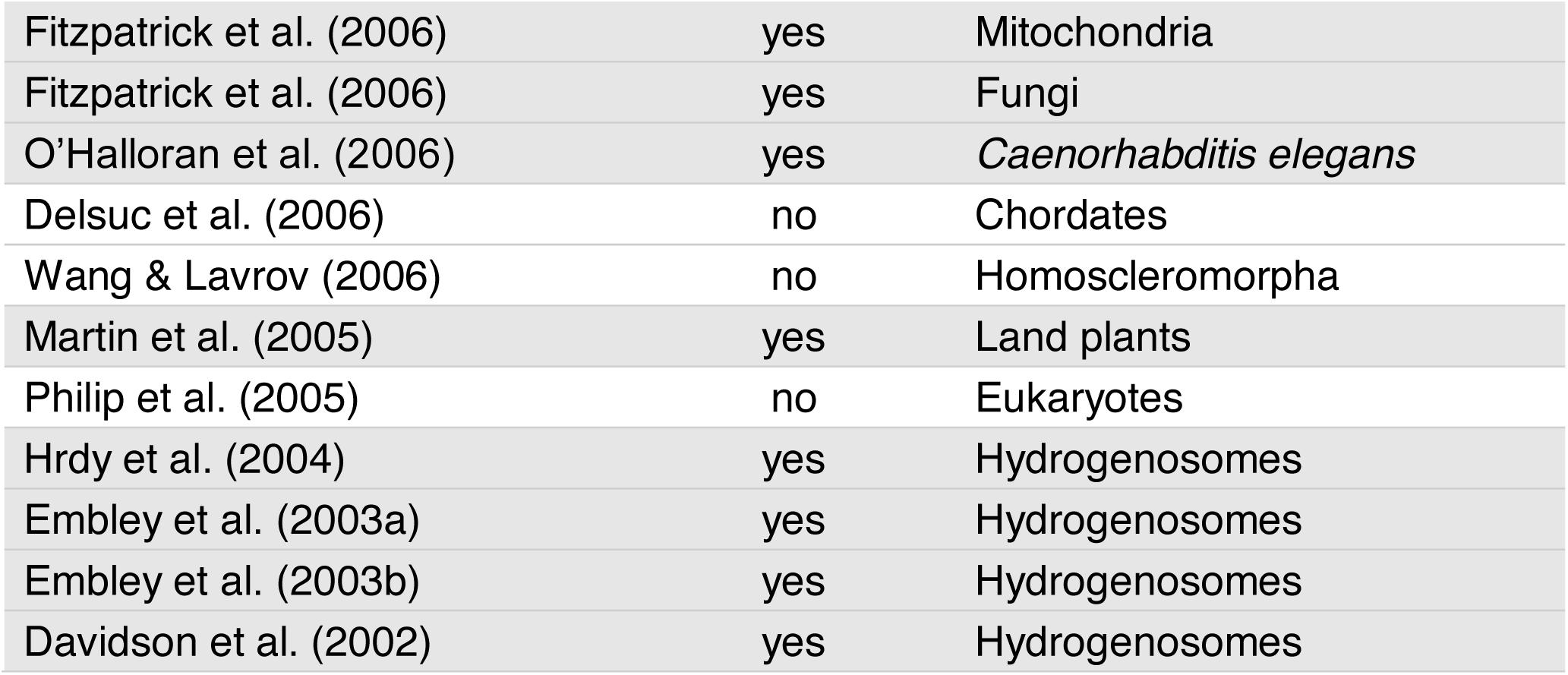
Publications that use 6-state amino acid recoding.

## Results

### The Efficacy of 6-state Recoding under a Compositional Heterogeneity Gradient

We simulated data with various levels of compositional heterogeneity by matching amino acid frequencies of four non-sister 5-taxa clades on a balanced 20-taxa tree and varying the length of the stem branches leading to those four clades (Figure 1A). We scored the ability of recoding and non-recoding approaches to recover the two compositionally heterogeneous 10-taxa clades (i.e., a clade containing all A and B taxa and a clade containing all C and D taxa). As compositional heterogeneity increased, the performance of the recoding approaches diminished at a slower rate than the non-recoding approaches (Figure 1B). However, in all cases tested, non-recoding approaches performed significantly better than recoding approaches, even under the highest levels of compositional heterogeneity and shortest stem branches (Table S2; Table S3).

**Figure 1.**
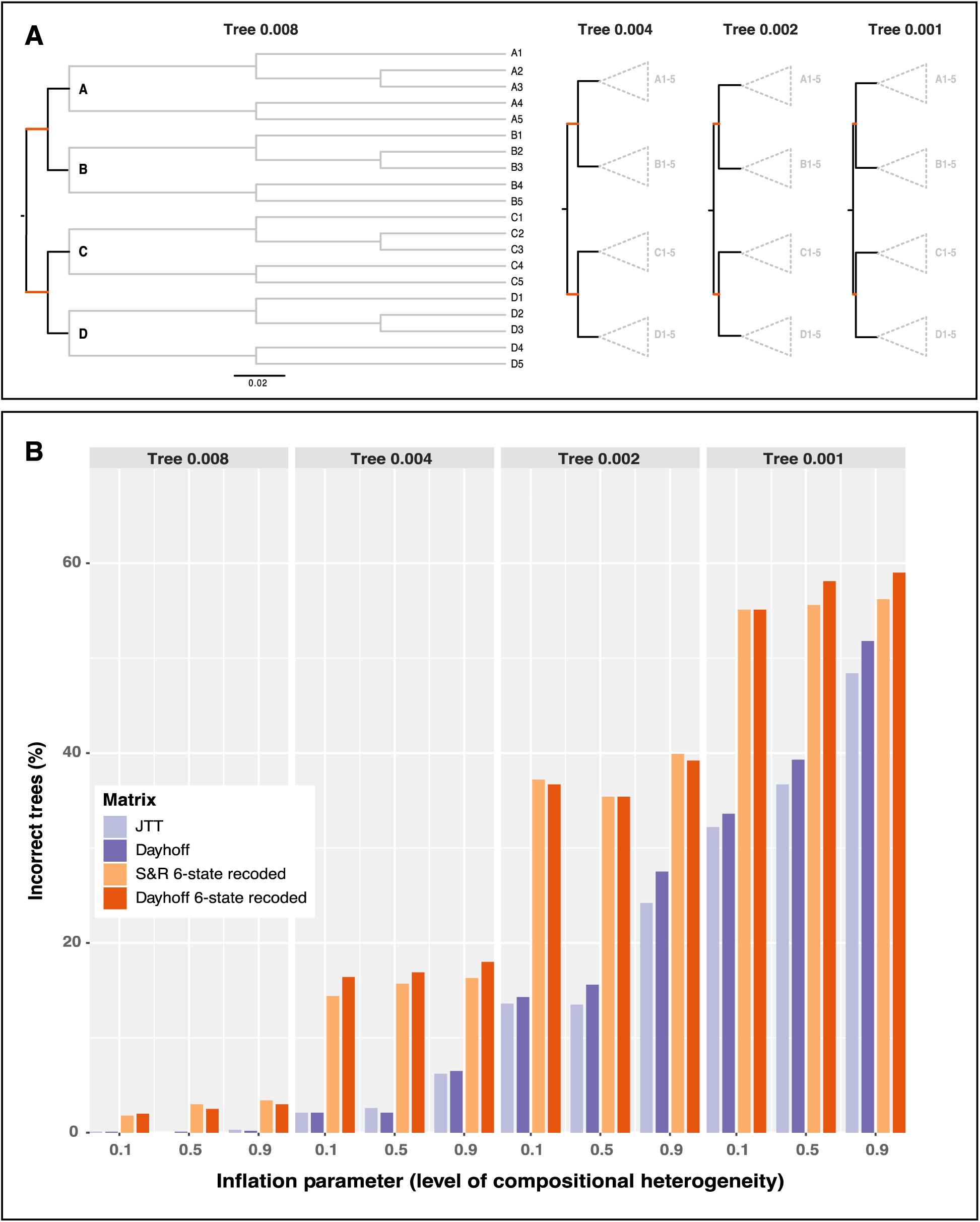
Six-state recoding approaches produce more incorrect trees under various levels of compositional heterogeneity. **(a)** Trees used for simulations. The value in the name of the tree (e.g., 0.008 in Tree 0.008) denotes the length in substitutions per site of the stem branches of the AB and CD clades (highlighted in orange). Decreasing the lengths of these branches increased the effect of compositional heterogeneity (Figure S3). **(b)** Percentage of 1000 trees that did not reconstruct a monophyletic group of taxa from clades A and B and monophyletic group of taxa from clades C and D.

### The Efficacy of 6-state Recoding under a Saturation Gradient

We simulated datasets on the Chang and Feuda trees under the Dayhoff and JTT models with increasing levels of saturation. Under all tested levels of saturation, phylogenetic reconstructions using the Dayhoff and LG models on non-recoded data matrices that were simulated under the Dayhoff model produced trees with fewer errors on average (as measured by Robinson-Foulds distances from the starting tree) than those that used the Dayhoff 6-state recoded matrix (Figure 2A). The results were similar for data simulated under the JTT model, where trees reconstructed with the JTT and LG models on non-recoded data matrices contained fewer errors on average across all tested levels of saturation compared to reconstructions with the S&R 6-state recoded matrix (Figure 2B). The results were consistent regardless of which topology (i.e., Chang or Feuda) was used for data simulations (Figure S2). As saturation increased, the performance of recoding approaches decreased at a faster rate than non-recoding approaches (Figure S2). T-tests performed for each branch length scaling factor parameter showed that Robinson-Foulds distances were significantly higher for recoded datasets compared to non-recoded datasets (p-value < 2.2e-16).

**Figure 2.**
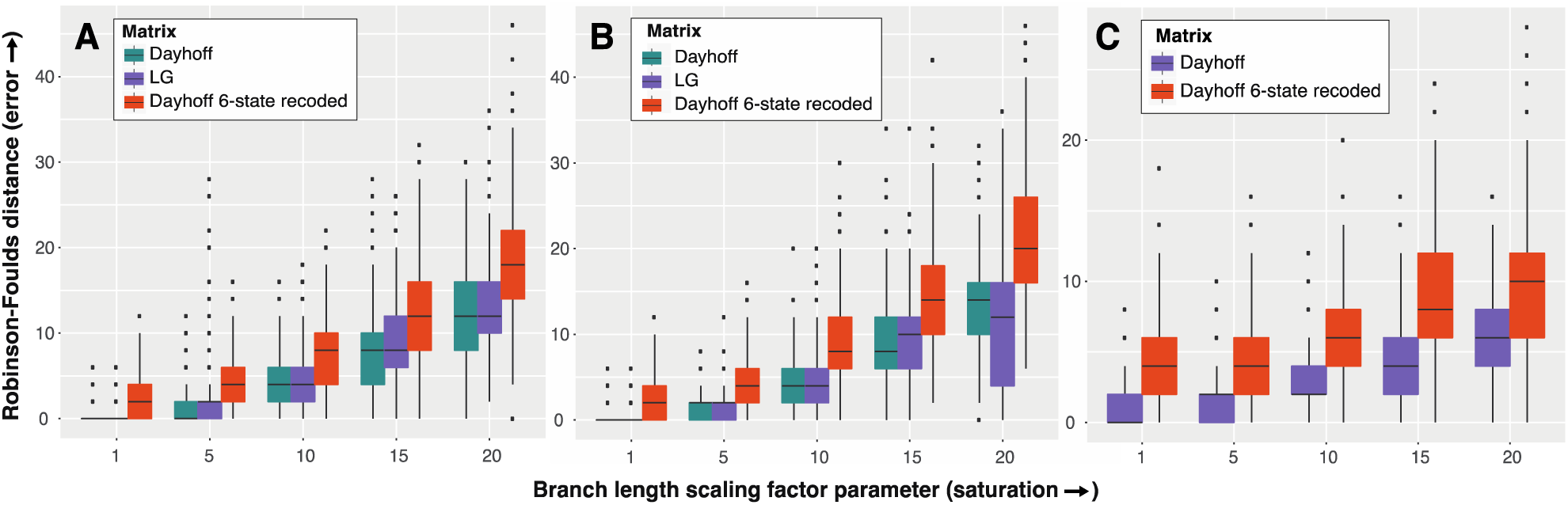
Six-state recoding approaches produce more errors under increasing levels of saturation. Robinson-Foulds distances of all 1,000 runs for each branch length scaling factor parameter. All data were simulated on the Chang tree topology. **(a)** Datasets simulated under the Dayhoff model. **(b)** Datasets simulated under the JTT model. **(c)** Datasets simulated under the GTR model using the amino acid rates of substitution, amino acid frequencies, and gamma rate heterogeneity estimated from the Chang dataset.

We also simulated data under the GTR model using the amino acid rates of substitution, amino acid frequencies, and gamma rate heterogeneity parameters estimated from the Chang dataset. Phylogenetic analyses of data simulated under GTR resulted in fewer errors on average when reconstructed with non-recoded Dayhoff matrices compared to reconstructions with the Dayhoff 6-state recoded matrices (Figure 2C). T-tests carried out for each branch length scaling factor parameter indicated that recoded approaches performed significantly worse than non-recoded approaches (p-value < 2.2e-16).

### The Effect of Alternative Recoding Strategies on Compositional Heterogeneity

We used the data simulated under inflation parameter 0.5 (mid-level of compositional heterogeneity) using the hypothetical tree 0.002 (short stem branches; Figure 1A) from the main compositional heterogeneity analysis to test Dayhoff 9-, 12-, 15-, and 18-state recoding strategies and compared the performance of these methods to Dayhoff 6-state recoding and non-recoding. As in the main compositional heterogeneity analysis outlined above, trees were assessed to determine if they recovered the two compositionally heterogeneous 10-taxa clades (i.e., AB and CD). The percentage of trees that passed these criteria increased as the number of Dayhoff states increased with Dayhoff 18-state recoding outperforming all other strategies including the non-recoding approach (Figure 3). Non-recoding outperformed all other recoding strategies except Dayhoff 12- and 15-state recoding under the highest level of compositional heterogeneity (inflation parameter 0.9; Figure 3C). Since the performance of Dayhoff-18 recoding surpassed the non-recoding method under all levels of compositional heterogeneity, we performed z-tests to determine if the differences in numbers of incorrect trees between analyses run with Dayhoff 18-state recoding and those run without recoding were significant. The difference was significant (p ≤ 0.05) only under the highest level of compositional heterogeneity (p-values for inflation parameters 0.1, 0.5, and 0.9: 0.2338, 0.1205, and 4.125e-06 respectively).

**Figure 3.**
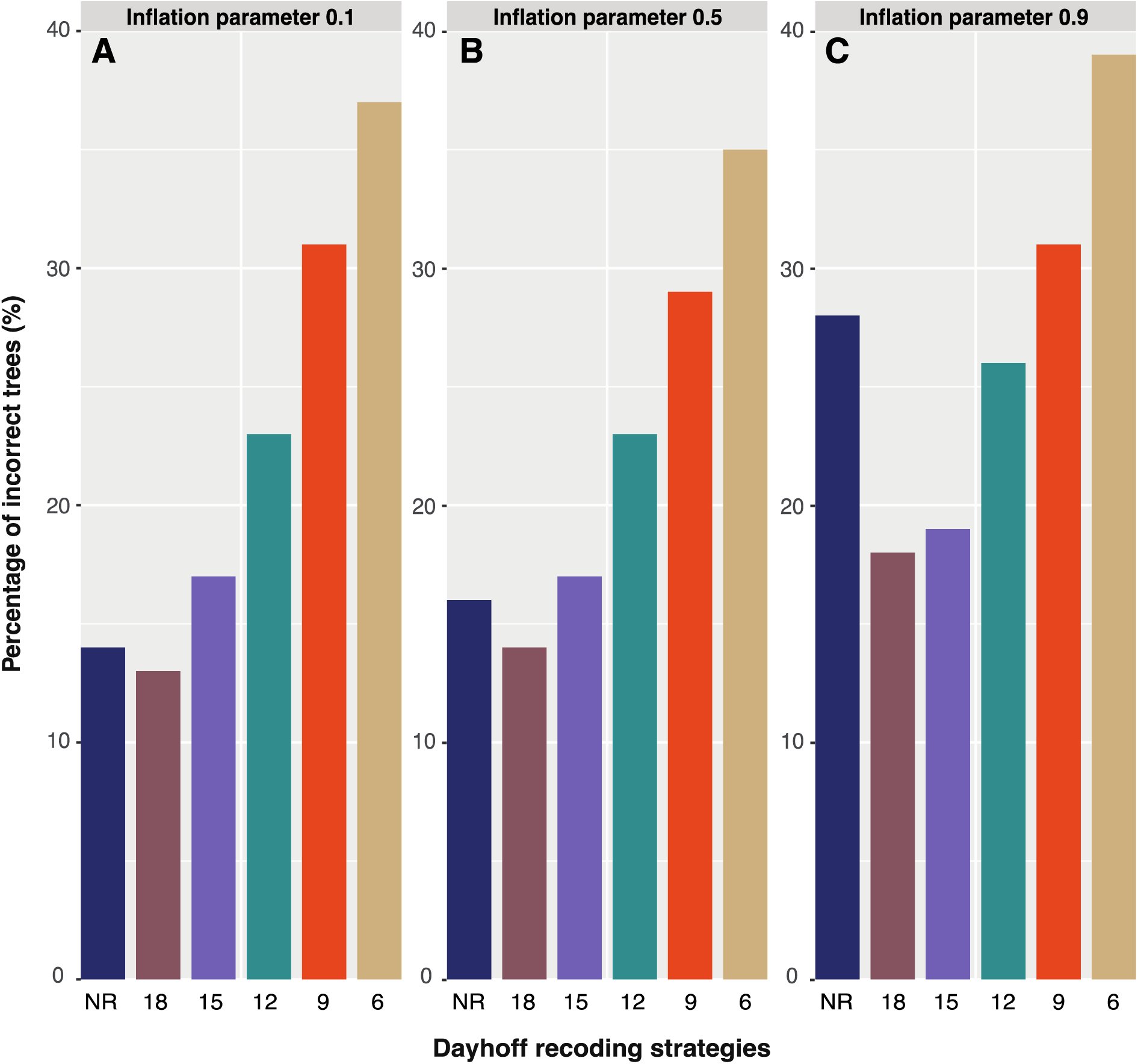
Dayhoff 9-, 12-, 15-, and 18-state recoding produce fewer incorrect trees than Dayhoff 6-state recoding under various levels of compositional heterogeneity. Trees were reconstructed by applying the non-recoded (NR) Dayhoff matrix or alternative Dayhoff recoding strategies (the number of states in the recoding strategy is indicated by digits). Incorrect trees did not include a monophyletic group of taxa from clades A and B and monophyletic group of taxa from clades C and D. The Y-axis refers to percentage out of 1,000 trees.

## Discussion

The philosophy underlying recoding strategies in phylogenetics is that sacrificing some information is beneficial in cases where homoplasy is high, as is the case when there is substantial heterogeneity in nucleotide or amino acid composition or when datasets are highly saturated. Six-state amino acid recoding has been proposed as a strategy to improve phylogenetic reconstruction in the presence of compositional heterogeneity and saturation (Embley et al. 2003; Hrdy et al. 2004; Martin et al. 2005). While there have been simulation analyses that compare different binning schemes (Susko and Roger 2007; Nesnidal et al. 2010), there are few if any studies that compare the accuracy of 6-state recoding to non-recoding approaches. In this study, we used simulations under gradients of compositional heterogeneity and saturation to compare the performance of 6-state amino acid recoding strategies. Remarkably, we found that non-recoding approaches outperformed 6-state recoding approaches in all of our comparisons. Our results show that while 6-state recoding seems to be less affected by increases in compositional heterogeneity, it does not overcome the penalty of information loss even under the highest levels of compositional heterogeneity (Figure 1B). Further, we found that 6-state recoding performs poorly when applied to highly saturated datasets. As such, we conclude that the costs of information loss associated with the 6-state recoding schemes are too great to justify applying these strategies.

It is possible that not all recoding strategies are inappropriate. Specifically, we found that our Dayhoff 9-, 12-, 15-, and 18-state recoding strategies performed better than the standard Dayhoff 6-state recoding approach for all tested levels of compositional heterogeneity (Figure 3). Dayhoff-18 recoding performed the best under all gradients of compositional heterogeneity and may comprise the optimum balance of minimizing compositional heterogeneity while maximizing information retention. However, we do not advocate blindly applying Dayhoff 18-state recoding, especially since significant (p 0.05) improvement only occurs under the most extreme compositional heterogeneity setting (0.9), which in no way reflects a realistic level. Instead we suggest that further simulation experiments with challenging topologies and realistic datasets are needed before adopting any amino acid recoding approach.

Applying a recoding method that is dataset specific may be another tactic to handle compositional heterogeneity or saturation. Susko and Roger (2007) and Nesnidal et al. (2010) applied this strategy by testing several recoding binning schemes informed by their datasets of interest. Tailoring the level and/or type of recoding to the amount of compositional heterogeneity and saturation, perhaps on a column-by-column basis, may be a successful approach, but further testing using such a tailored method would be necessary. Since only a handful of studies have investigated different recoding schemes, it is clear that more analyses are required to gain an understanding of the impact of alternative recoding methods for compositionally heterogeneous and/or saturated datasets.

### Implications

There are at least 77 publications that use 6-state amino acid recoding, with the first seven months of 2019 seeing more than any year to date (Table 1). Many of these studies have proposed controversial topologies with profound implications across the tree of life including bacteria, archaea, unicellular eukaryotes, fungi, animals, and plants. We have shown that 6-state recoding greatly reduces information content and therefore often results in suboptimal phylogenetic reconstructions. We therefore advocate caution when interpreting results stemming from analyses that have employed 6-state recoding and contend that publications in which 6-state recoding analyses had a substantial effect on the conclusions be revisited.

## Materials and Methods

### Reproducibility and Transparency Statement

Custom scripts, command lines, and data used in these analyses are available at https://github.com/josephryan/Hernandez_Ryan_2019_Recoding Sim

To maximize transparency and minimize confirmation bias, all analyses were pre-planned using phylotocol (DeBiasse and Ryan 2018) and pre-registered using the Center for Open Science’s pre-registration platform (https://osf.io/smj6k/ and https://osf.io/6ubgj/). We made three changes to our original plan during the life of this project, and these changes were documented and justified in the phylotocol available on our GitHub repository (URL above).

### Overview of Empirical Datasets Employed

The following methods can be divided into two main analyses: compositional heterogeneity and saturation. Both analyses employ empirical data from the following papers: Chang et al. (2015) hereafter “Chang,” and Feuda et al. (2017) hereafter “Feuda.” The topologies from Chang and Feuda are based on the same dataset which is made up of 51,940 amino acid positions from 77 taxa representing a wide range of animals and 9 non-animal outgroups. Feuda extensively applied 6-state amino acid recoding to this dataset in a reanalysis of the Chang study, which did not use recoding.

For the compositional heterogeneity analysis, we use several hypothetical 20-taxa symmetrical trees which consist of 4 clades (named A, B, C, and D) made up of 5 taxa each (Figure 1A), and apply global parameters estimated from the Chang dataset. For the saturation analysis, we use the topologies reported in Chang and Feuda. More details on these analyses are provided below.

### Testing 6-state Recoding Performance on Compositional Heterogeneity

We used the script comphet.pl (available in our GitHub repository) to simulate amino acid data in P4 (Foster 2004) on four hypothetical 20-taxa balanced trees (Figure 1A). We simulated sequences that were 1,000 amino acids in length under the GTR model using the amino acid rates of substitution from the Chang dataset. To introduce compositional heterogeneity, we generated one set of amino acid frequencies for clades A and C and a different set of frequencies for clades B and D. For clades B and D, we used the amino acid frequencies from the Chang dataset. For clades A and C, frequencies for the following pairs of amino acids ((A,L), (R,K), (N,M), (D,F), (C,P), (Q,S), (E,T), (G,W), (H,Y), (I,V); chosen based on an alphabetical pattern) were determined by adjusting each frequency by X, where X is the inflation parameter (i.e., 0.1, 0.5, 0.9) multiplied by the lowest frequency of the pair. The amino acid of the pair with the lowest frequency is incremented by X and the other is decremented by X.

For example, the Chang frequencies for the amino acids R and K are 0.063 and 0.080 respectively. These frequencies were used for clades B and D without adjustment. To determine the increment value X under the inflation parameter 0.1, we multiplied the frequency of R, which is the lowest of the pair, by 0.1 (X=0.0063). We then added X to the Chang frequency of R (0.063 + 0.0063) and subtracted X from the Chang frequency of K (0.080 − 0.0063). We rounded these values to 3 decimal places (to work with P4) for a final set of frequencies of R = 0.069 and K=0.074.

Using this algorithm to generate frequencies, we performed 1,000 simulations for each combination of the four hypothetical 20-taxa trees and the three inflation parameters (i.e., 0.1, 0.5, and 0.9) resulting in 12,000 total datasets. To verify that these datasets displayed compositional heterogeneity between clades A and C compared to clades B and D, we computed amino acid frequencies on the simulated datasets and summed the difference in frequencies across all replicates. We subtracted the average difference in amino acid frequencies between clades A and C and between clades B and D (clade pairs with homogeneous composition) from the average difference in amino acid frequencies between clades A and B, A and D, B and C, and C and D (clade pairs with heterogenous composition) to generate a compositional heterogeneity (comp-het) index value. Datasets with comp-het index values closer to 0 are characterized by low compositional heterogeneity, while datasets with higher comp-het index values are characterized by high compositional heterogeneity.

We recoded each simulated dataset with both Dayhoff 6-state recoding and S&R 6-state recoding, and then reconstructed maximum-likelihood trees of the recoded datasets using the GTR multi-state model and of the non-recoded datasets using the Dayhoff and JTT models in RAxML (Stamatakis 2014). In total we produced 48,000 phylogenies for testing compositional heterogeneity. We used the script is_mono.pl (available in our GitHub repository) to determine whether each tree recovered a monophyletic group that included all A and B taxa, which by definition would include a monophyletic group that included all C and D taxa. We did not test for more fine-scale relationships as our goal was to evaluate the degree to which the applied level of compositional heterogeneity was pulling together the compositionally homogenous clades A and C, and B and D. We calculated the percentage of incorrect trees using the above criteria for each combination of model, recoding type (including no recoding), and level of applied compositional heterogeneity (i.e., inflation parameter), and performed a z-test to compare the proportions of incorrect trees between non-recoding and recoding approaches.

### Testing 6-state Recoding Performance on Saturation

We used Seq-Gen (Rambaut and Grass 1997) to simulate the evolution of amino acids on the Chang and Feuda topologies. First, we confirmed that increasing the branch length scaling factor parameter in Seq-Gen linearly increased levels of saturation (Figure S1) using the script seq-gen_saturation_test.pl (available in the accompanying GitHub repository). Next, we performed 1,000 simulations per combination of tree topology (Chang and Feuda), branch length scaling factor parameter (1– 20), and model of amino acid substitution (either Dayhoff or JTT) for a total of 80,000 datasets. We simulated an additional 1,000 datasets on the Chang topology for a subset of branch length scaling factor parameters (1, 5, 10, 15, 20) under the GTR model using the amino acid rates of substitution, amino acid frequencies, and gamma rate heterogeneity from the Chang dataset, bringing the grand total to 85,000 datasets. Each dataset included 1,000 amino acid columns.

For simulations performed on the Chang topology, we increased the branch length scaling factor parameter from 1 to 20 in increments of 1. The Feuda topology was produced from recoding the Chang dataset (Feuda et al. 2017), and because trees produced from recoded data have substantially fewer substitutions and therefore shorter branch lengths, we incremented branch lengths by a factor of 2.6 for the Feuda tree (based on our calculation that the sum of branch lengths in the recoded tree was 2.6 shorter than the sum of branch lengths in the non-recoded Chang tree).

We performed maximum-likelihood analyses with RAxML for each set of sequences produced from simulations over the Chang and Feuda topologies. For the datasets simulated with Dayhoff and JTT substitution models, we reconstructed trees using the generating model, the 6-state recoding scheme derived from that model, and for a subset of branch length scaling factor parameters (1, 5, 10, 15, 20) we also reconstructed trees using LG, a sub-optimal model in this context, as it was not the model used for the simulations. For the datasets simulated with the GTR substitution model, we generated trees using Dayhoff and Dayhoff 6-state recoding. We produced 180,000 phylogenies in total to test saturation. To test the performance of each recoding (or non-recoding) scheme, we used TOPD/FMTS (Puigbo et al. 2007) to calculate Robinson-Foulds distances (Robinson and Foulds 1981) between the topology used for simulation (i.e., Chang or Feuda) and the reconstructed trees generated from simulated sequences. We used a t-test to determine if there were significant differences in Robinson-Foulds distances between recoded and non-recoded datasets for each branch length scaling factor.

### Testing Alternative Recoding Strategies on Compositional Heterogeneity

To test the effect of number of states on recoding strategies, we developed alternative Dayhoff 9-, 12-, 15-, and 18-state recoding strategies. The first step in these analyses was to determine the optimal amino acid binning strategy for each number of tested states. Since the number of possible bins for each state is finite, ideally, we would use an exhaustive algorithm to identify the binning scheme that maximizes the sum of intra-bin substitution scores using the Dayhoff matrix. Unfortunately, as pointed out by Susko and Roger (2007), the number of possible bins is very large (e.g., there are roughly 1.5 × 1013 choices of bins under an 8-state recoding strategy) and an exhaustive algorithm is computationally intractable. Instead, we generated scores (see score.pl in our GitHub repository) for several binning schemes that incorporated subsets of the Dayhoff 6-state recoding bins and chose the best-scoring binning strategies from this set (Table S1). We also compared our best binning strategies to those proposed in Susko and Roger (2007) and in all cases, the scores we generated were higher, except for one which had an equal score (not entirely surprising given that the Susko and Roger bins were optimized for JTT recoding).

We compared the binning schemes that scored the highest for each recoding strategy (Table 2) against the Dayhoff and Dayhoff 6-state recoded matrices by testing their performance under reasonably high levels of compositional heterogeneity. We recoded the data that we simulated for the compositional heterogeneity analysis (data simulated with inflation parameter 0.5 using the hypothetical tree 0.002 (Figure 1A)) using our Dayhoff 9-, 12-, 15-, and 18-state recoding strategies and reconstructed maximum-likelihood trees in RAxML. As in the main compositional heterogeneity analysis outlined above, we used the script is_mono_comphet.pl to test if the ten taxa labeled A and B were monophyletic and likewise the ten taxa labeled C and D were monophyletic. We also performed a z-test to compare the proportion of incorrect trees produced under Dayhoff-18 recoding (see Results for rationale) to those produced under non-recoding.

**Table 2.** Best scoring binning schemes optimized on the Dayhoff matrix.

| Dayhoff recoding | Binning scheme |
| --- | --- |
| 9-state | DEHNQ ILMV FY AST KR G P C W |
| 12-state | DEQ MLIV FY KHR G A P S T N W C |
| 15-state | DEQ ML IV FY G A P S T N K H R W C |
| 18-state | ML FY I V G A P S T D E Q N H K R W C |

## Supporting information

Supplementary commands, parameters, and version numbers of programs used in this analysis

## Supplementary Material

All commands and versions of software used in this study are provided in the supplementary material. All data and scripts are available in the following GitHub repository: https://github.com/josephryan/Hernandez_Ryan_2019_Recoding Sim.

## Acknowledgements

We thank Melissa DeBiasse for providing comments on an earlier version of the manuscript and Gordon Burleigh, Christine Schnitzler, Marta Wayne, and Bryan Kolaczkowski for feedback on this project during A.M.H.’s committee meeting. The authors would like to express their thanks to David Swofford and Gavin Naylor for influential discussions at an early stage of the project. The views expressed in this paper do not necessarily reflect the views of those acknowledged.

