## Supplementary figures and tables for "Six-state amino acid recoding is not an effective strategy to offset the effects of compositional heterogeneity and saturation in phylogenetic analyses"

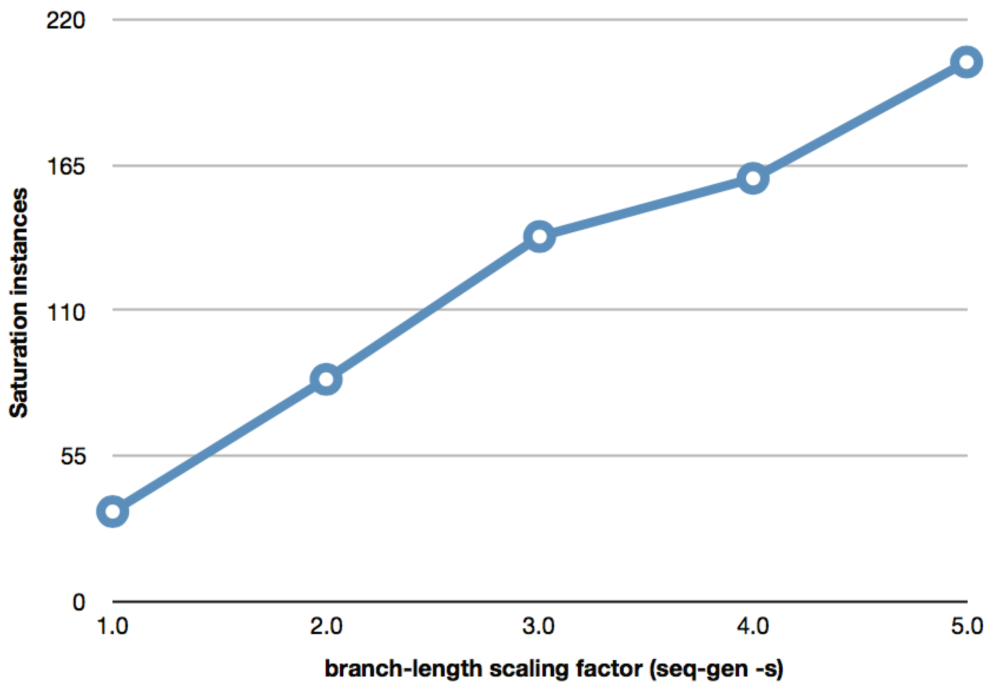


**Figure S1. Relationship between the branch length scaling factor and saturation.** Saturation instances are the number of times an amino acid position changed from one amino acid to another and back. The Y axis is an underestimate since only changes at internal nodes are considered.

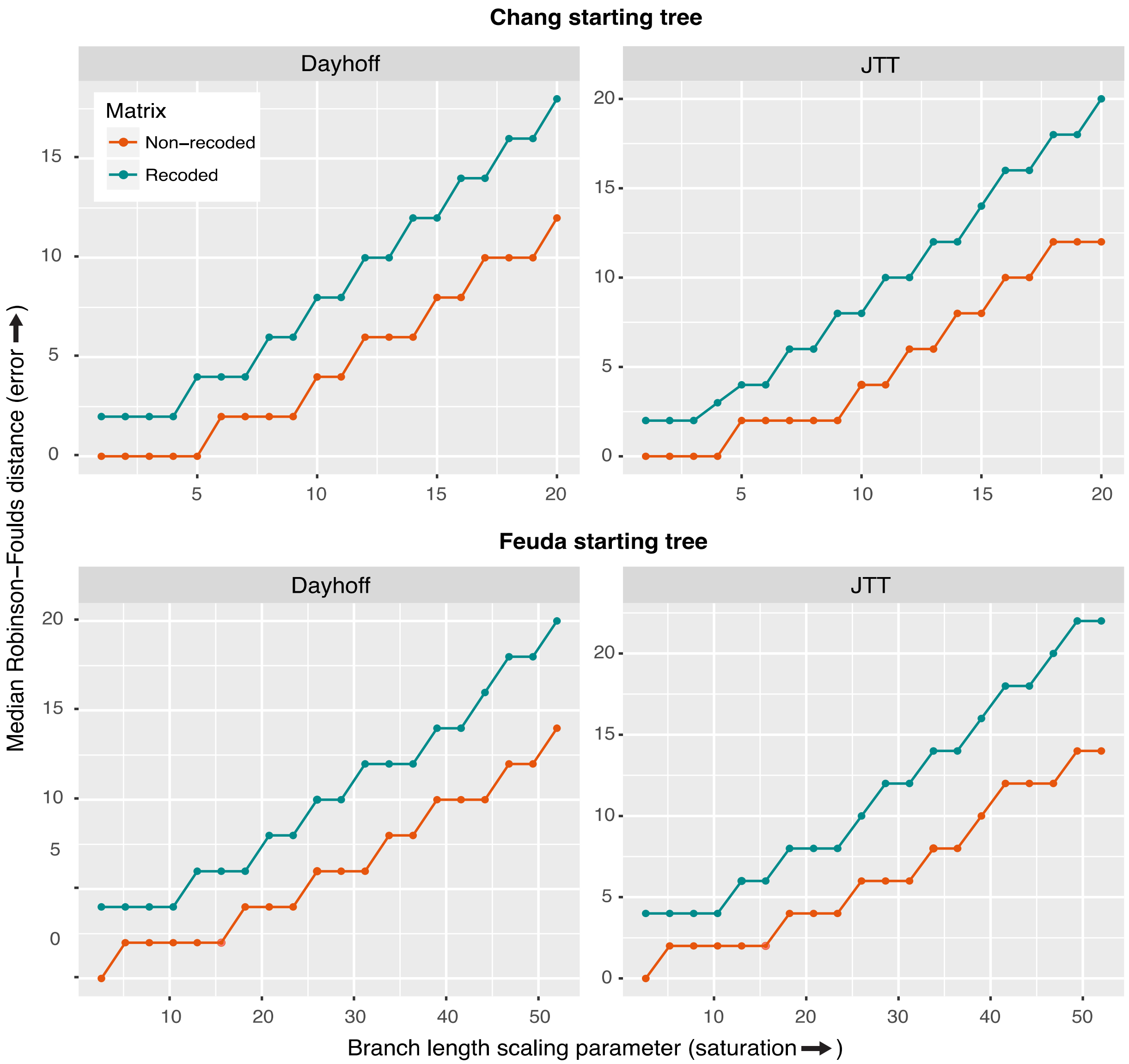


**Figure S2. Median Robinson-Foulds distances for non-recoded and recoded datasets across a gradient of saturation levels.** Datasets simulated over: **(a)** Chang topology using the Dayhoff model. **(b)** Chang topology under the JTT model. **(c)** Feuda topology under the Dayhoff model. **(d)** Feuda topology using the JTT model.

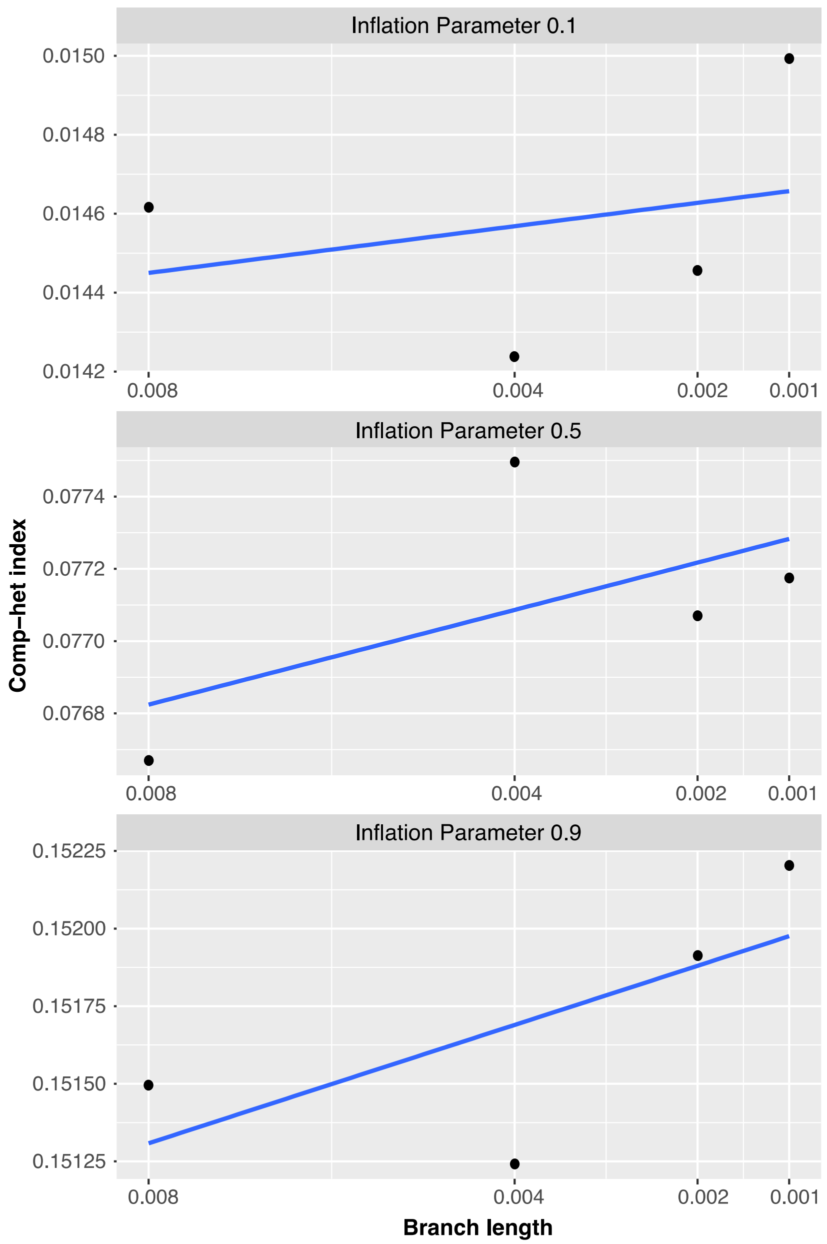


**Figure S3. Decreases in the length of stem branches of the AB and CD clades (highlighted in red in Figure 1A) increase compositional heterogeneity.**

| Dayhoff recoding | Binning scheme | Score |
|---|---|---|
| 9-state | **DEHNQ ILMV FY AST KR G P C W** | **47** |
| | EHNQ IMTV FY ASL KR G P C W | 23 |
| | ILMV DEQ AGPST KN FY H C W R | 37 |
| | ILMVT DENQ APS FY KR H G C W | 37 |
| 12-state | **DEQ MLIV FY G A P S T N KHR W C** | **35** |
| | DEQ M LIV FY G APST N K H R W C | 27 |
| | DEQ M LIV FY GAPS T N K H R W C | 26 |
| | DEQ M LIV FY GAPT S N K H R W C | 24 |
| | DEQ MLIV FY GAP T S N K H R W C | 31 |
| | D E Q MLIV FY GAP S T N KHR W C | 29 |
| 15-state | **DEQ ML IV FY G A P S T N K H R W C** | **22** |
| | ML IV G A P S T DE Q N K H RFY W C | 10 |
| | ML IV G A P S T DE Q N K H R FYW C | 18 |
| | ML IV G A P S T DEQ N K H R FY W C | 22 |
| | ML IV G A P S T DEN Q K H R FY W C | 21 |
| | ML I V GAP S T DE Q N K H R FY W C | 15 |
| | ML IVG A P S T DE Q N K H R FY W C | 14 |
| | ML IV G A P S T DEN Q K H R FY W C | 21 |
| | ML IV G A P S T D E QN KHR FY W C | 20 |
| | ML IV G A P S T DE Q N KHR FY W C | 19 |
| 18-state | IV ML G A P S T D E Q N H K R F Y W C | 8 |
| | IV M L G A P S T D E Q N H K R F Y WC | -4 |
| | IV M L G A P S T DE Q N H K R F Y W C | 7 |
| | IV M L G A P S T D E QN H K R F Y W C | 5 |
| | IV ML G A P S T D E Q N H K R F Y W C | 8 |
| | MV IL G A P S T D E Q N H K R F Y W C | 4 |
| | MI VL G A P S T D E Q N H K R F Y W C | 4 |
| | ML IV G A P S T D E Q N H K R F Y W C | 8 |
| | ML I V GA P S T D E Q N H K R F Y W C | 5 |
| | ML I V G AP S T D E Q N H K R F Y W C | 5 |
| | ML I V G A P ST D E Q N H K R F Y W C | 5 |
| | ML I V G A P S T DE Q N H K R F Y W C | 7 |
| | ML I V G A P S T D E QN H K R F Y W C | 5 |
| | ML I V G A P S T D EQ N H K R F Y W C | 6 |
| | ML I V G A P S T D E Q N H K RF Y W C | 0 |
| | **ML I V G A P S T D E Q N H K R FY W C** | **11** |
| | ML I V G A P S T D E Q N HK R F Y W C | 4 |
| | ML I V G A P S T D E Q N H KR F Y W C | 7 |
| | ML I V G A P S T D E Q N K HR F Y W C | 6 |

**Table S1. Binning schemes tested for optimization of scores based on the Dayhoff matrix.** The best scoring schemes are bolded.

| Tree | Inflation parameter | Non-recoded (% error) | Recoded (% error) | P-value |
|---|---|---|---|---|
| 0.008 | 0.1 | 0.1 | 2.0 | 4.28E-05 |
| | 0.5 | 0.1 | 2.5 | 3.23E-06 |
| | 0.9 | 0.2 | 3.0 | 9.08E-07 |
| 0.004 | 0.1 | 2.1 | 16.4 | < 2.2e-16 |
| | 0.5 | 2.1 | 16.9 | < 2.2e-16 |
| | 0.9 | 6.5 | 18.0 | 1.63E-13 |
| 0.002 | 0.1 | 14.3 | 36.7 | < 2.2e-16 |
| | 0.5 | 15.6 | 35.4 | < 2.2e-16 |
| | 0.9 | 27.5 | 39.2 | 3.54E-06 |
| 0.001 | 0.1 | 33.6 | 55.1 | 3.35E-13 |
| | 0.5 | 39.3 | 58.1 | 1.04E-09 |
| | 0.9 | 51.8 | 59.0 | 0.01646 |

**Table S2. Z-test for significant differences between percentages of error for trees solved with Dayhoff and Dayhoff 6-state recoding.**

| Tree | Inflation parameter | Non-recoded (% error) | Recoded (% error) | P-value |
|---|---|---|---|---|
| 0.008 | 0.1 | 0.1 | 1.8 | 0.000121 |
| | 0.5 | 0.0 | 3.0 | 5.96E-08 |
| | 0.9 | 0.3 | 3.4 | 4.07E-07 |
| 0.004 | 0.1 | 2.1 | 14.4 | < 2.2e-16 |
| | 0.5 | 2.6 | 15.7 | < 2.2e-16 |
| | 0.9 | 6.2 | 16.3 | 1.31E-11 |
| 0.002 | 0.1 | 13.6 | 37.2 | < 2.2e-16 |
| | 0.5 | 13.5 | 35.4 | < 2.2e-16 |
| | 0.9 | 24.2 | 39.9 | 3.60E-10 |
| 0.001 | 0.1 | 32.2 | 55.1 | 5.97E-15 |
| | 0.5 | 36.7 | 55.6 | 3.05E-10 |
| | 0.9 | 48.4 | 56.2 | 0.008637 |

**Table S3. Z-test for significant differences between percentages of error for trees solved with JTT and S&R 6-state recoding.**
