## Supplementary commands, parameters, and version numbers of programs used in this analysis for "Six-state amino acid recoding is not an effective strategy to offset the effects of compositional heterogeneity and saturation in phylogenetic analyses"

**Commands, parameters, and version numbers of programs to reproduce data simulation and phylogenetic analyses**

### 1. Setting up the environment

#### 1.1 Clone the GitHub repository for this paper and set the REPO variable.

```
git clone https://github.com/josephryan/Hernandez_Ryan_2019_RecodingSim
```

```
cd Hernandez_Ryan_2019_RecodingSim
```

```
export REPO=`pwd`
```

#### 1.2 Edit Servers.pm

The following commands run on a single modern processor would take more than 6 years to complete. We used 4 servers with more than 300 CPUs between them. To distribute jobs between servers, we developed scripts that will generate individual shell scripts. We provide commands to run these individual scripts on different servers. To distribute the jobs on multiple servers (or multiple CPUs on a single server), the following file must be edited.

$REPO/00-MODULES/JFR/Servers.pm

**The following external programs are expected to be installed and in $PATH. The versions applied to our analyses are listed in parentheses.**

Python (version 2.7.16)
P4 (version 1.2.0)
ete3 (version 3.1.1)
six (version 1.12.0)
RAxML (version 8.2.11)
Seq-Gen (version 1.3.2)
TOPD/FMTS (version 4.6)
R (version 3.3.1) - ggplot2 package (version 2.2.1)
RStudio (version 1.2.1335) - ggplot2 package (version 3.1.1)

### 2. Testing Compositional Heterogeneity

#### 2.1 Simulate datasets for null distribution (no induced compositional heterogeneity)

**2.2.1** In the directory 02-COMPOSITIONAL_HETEROGENEITY/01-NULL_DISTRIBUTION/01-TREE0008, make the directory ‘scripts’ and run the following command to generate several shell scripts that perform 1,000,000 simulations.

```
mkdir scripts
```

```
perl split_random_runs.pl TREE0008
```

**2.2.2** The following command will execute the shell scripts to simulate amino acid data on hypothetical tree 0.008 (Figure 1A) under the GTR model with amino acid frequencies and transition rates estimated from the Chang dataset. Shell scripts will generate simulated sequences in PHYLIP format within each output directory, p4 scripts (p4.x) that were used to perform simulations, and ".out" files with comp-het index values for each simulated dataset.

```
ls -1 scripts | grep 'myserver*' | perl -ne 'chomp; print "sh scripts/$_ &\n”;’ | sh
```

**2.2.3** Concatenate all ".out" files. The comp-het index values in these files are used as a null distribution to statistically compare to the comp-het indices from datasets that were simulated under gradients of compositional heterogeneity.

```
cat *.out >> all.out
```

**2.3 Simulate compositionally heterogeneous datasets**

**2.3.1** In the directory 02-COMPOSITIONAL_HETEROGENEITY/02-HETEROGENEOUS_DATA, run the following commands to produce datasets with increasing compositional heterogeneity. The script comphet.pl prints out the comp-het index value for the simulated dataset and its associated p-value. The p-value is calculated by comparing the comp-het index of the simulated dataset to the null distribution of comp-het indices.

```
cd 01-TREE0008.1

perl comphet.pl TREE0008 0.1 all.out > pvals.0.1.out

cd 02-TREE0008.5

perl comphet.pl TREE0008 0.5 all.out > pvals.0.5.out

cd 03-TREE0008.9

perl comphet.pl TREE0008 0.9 all.out > pvals.0.9.out
```

**2.4 Relationship between length of the stem branches of the AB and CD clades and compositional heterogeneity**

We looked at the relationship between the lengths of the stem branches of the AB and CD clades for each tree (highlighted in red in Fig. 1A) and the comp-het index values to determine how branch lengths were impacting compositional heterogeneity.

**2.4.1** Run the script comphet_index_scatter.R in RStudio to produce a scatter plot showing the relationship between the length of the stem branches of the AB and CD clades for each tree and comp-het index values for each inflation parameter tested.

**2.5 Recode datasets and create maximum-likelihood analysis shell scripts**

**2.5.1** In each of the directories 01-TREE0008.1, 02-TREE0008.5, and 03-TREE0008.9 make the directory 'scripts' and run the script chunkify_comphet_trees.pl to recode the compositionally heterogeneous dataset using Dayhoff 6-state recoding and S&R 6-state recoding. This script also generates shell scripts to perform maximum-likelihood analyses in RAxML.

```
mkdir scripts
```

```
perl chunkify_comphet_trees.pl
```

#### 2.6 Phylogenetic reconstructions

**2.6.1** Run the following command in each of the directories 01-TREE0008.1, 02-TREE0008.5, and 03-TREE0008.9 to execute the shell scripts generated from chunkify_comphet_trees.pl to perform maximum-likelihood analyses on both recoded and non-recoded datasets. Recoded datasets were reconstructed under the MULTIGAMMA multi-state model with GTR and non-recoded datasets were reconstructed under the Dayhoff and JTT models.

```
ls -1 scripts/ | grep 'myserver*' | perl -ne 'chomp; print "sh
scripts/$_&\n";' | sh
```

#### 2.7 Score trees

**2.7.1** Run the following command in each of the directories 01-TREE0008.1, 02-TREE0008.5, and 03-TREE0008.9 to determine the proportion of trees that do not correctly reconstruct a monophyletic group including taxa from clades A and B and a monophyletic group including taxa from clades C and D. The output file identifies which trees were incorrectly reconstructed and indicates the proportion of incorrect trees produced under each method (recoding and non-recoding).

```
perl is_mono.pl DAYHOFF > is_mono_dayhoff.out
```

```
perl is_mono.pl JTT > is_mono_jtt.out
```

#### 2.8. Perform 2.1 – 2.7 using hypothetical trees 0.004, 0.002, and 0.001 (Figure 1A).

**2.8.1** Repeat analyses 2.1-2.7 replacing TREE0008 with TREE0004, TREE0002, and TREE0001.

#### 2.9 Bar graph and z-test for percentage of incorrect trees

**2.9.1** Run the script comphet_bargraph.R in RStudio to generate a bar graph showing the percentage of incorrect trees reconstructed under each method, tree used for simulation, and level of compositional heterogeneity tested.

**2.9.2.** Run the script ztest_proportions_comphet.R in RStudio to perform z-tests on the proportions of incorrect trees reconstructed under recoding and non-recoding methods for each tree used for simulation and level of compositional heterogeneity tested.

### 3. Testing Saturation

#### 3.1 Testing the association between the branch length scaling factor parameter and saturation

We performed a simulation experiment with a six-taxa bifurcating tree (test.tre) and ran five instances of Seq-Gen with incrementing “branch length scaling factor” parameters (i.e., 1.0, 2.0, 3.0, 4.0, and 5.0) and the "write ancestral sequences for each node" parameter (i.e., -w a) set, which printed ancestral sequences at each node. We then calculated the number of saturation events (i.e., the number of times a change in an amino acid led to the appearance of an amino acid that had been in that same position in a prior ancestral node) using the script seq-gen_saturation_test.pl.

**3.1.1** In the directory 03-SATURATION/01-SATURATION_TEST, run seq-gen_saturation_test.pl to perform the simulations outlined above.

```
perl seq-gen_saturation_test.pl
```

**3.2 Simulations**

**3.2.1** In the directory 03-SATURATION/02-SEQ_GEN_CHANG/01-DAYHOFF, use the following shell scripts to reproduce the simulations we performed in Seq-Gen using the Chang topology, the PAM250 (Dayhoff) model, and the branch length scaling factor parameters described in the manuscript. These scripts will generate a total of 20,000 datasets, create new directories for each topology/model/branch length scaling factor parameter tested, and parse the datasets into separate PHYLIP files within their corresponding directories.

```
sh run_seqgen_pam_chang.sh
```

```
sh divide_chang_pam.sh
```

**3.2.2** Run the following shell scripts in the directory 03-SATURATION/02-SEQ_GEN_CHANG/02-JTT to perform equivalent simulations under the JTT model.

```
sh run_seqgen_jtt_chang.sh
```

```
sh divide_chang_jtt.sh
```

**3.2.3** The following commands and shell scripts reproduce the simulations we performed using the Feuda topology, the PAM250 model, and the branch length scaling factor parameters described in the manuscript. These scripts will generate a total of 20,000 datasets, create new directories, and parse data as in 3.2.1. Run these commands in the directory 03-SATURATION/03-SEQ_GEN_FEUDA/01-DAYHOFF.

```
sh run_seqgen_pam_feuda.sh
```

```
sh divide_feuda_pam.sh
```

**3.2.4** Running the following shell scripts in the directory 03-SATURATION/03-SEQ_GEN_FEUDA/02-JTT performs equivalent simulations using the Feuda topology and the JTT model.

```
sh run_seqgen_jtt_feuda.sh
```

```
sh divide_feuda_jtt.sh
```

**3.3 Recode datasets**

**3.3.1** In the directory 03-SATURATION, create a new directory called ‘scripts.’ Run chunkify.pl to convert simulated datasets to Dayhoff 6-state recoded and S&R 6-state recoded datasets and generate shell scripts to perform maximum-likelihood analyses in RAxML.

```
mkdir scripts
```

```
perl chunkify.pl
```

#### 3.4 Phylogenetic reconstructions

**3.4.1** Execute the scripts generated from chunkify.pl to perform maximum-likelihood analyses on both recoded and non-recoded datasets. Recoded datasets were reconstructed under the MULTIGAMMA multi-state model with GTR and non-recoded datasets were reconstructed under the Dayhoff model.

```
ls -1 scripts | grep 'myserver*’ | perl -ne 'chomp; print "sh scripts/$_
&\n";' | sh
```

#### 3.5. Score trees

**3.5.1** Run the following script to calculate Robinson-Foulds distance (RFD) values between the tree used for simulation and the recoded and non-recoded trees for each branch length scaling factor parameter tested. The script will also perform t-tests in R to determine if there are significant differences in RFDs between recoded and non-recoded trees, as well as generate box-plots to visualize the data.

```
perl compare_trees_to_sim.pl
```

#### 3.6 Line graph of median RFDs for non-recoded and recoded datasets

**2.6.1** Run the script Robinson_Foulds_medians_fig.R in RStudio to generate a line graph displaying median Robinson-Foulds distances for each tree, model, and branch length scaling factor parameter tested.

#### 3.7 Phylogenetic reconstructions under the LG model

**3.7.1** Datasets simulated over the Chang tree with branch length scaling factors 1, 5, 10, 15, and 20 were reconstructed using the LG model. To reproduce this analysis, in the directory 03-SATURATION make a new directory called ‘LG_scripts’. Run the script chunkify_LG.pl to generate shell scripts for phylogenetic reconstructions in RAxML.

```
mkdir LG_scripts
```

```
perl chunkify_LG.pl
```

**3.7.2** Use the following command to execute the shell scripts generated from chunkify_LG.pl that perform maximum-likelihood analyses using the LG model.

```
ls -1 scripts | grep 'myserver*' | perl -ne 'chomp; print "sh scripts/$_
```

```
&\n";' | sh
```

**3.8 Scoring and comparing trees reconstructed with the LG model**

**3.8.1** Run compare_trees_to_sim_LG.pl to score trees reconstructed under the LG model using the RFD in TOPD/FMTS. This script performs t-tests to compare the RFD values between the trees reconstructed with LG and those reconstructed with 6-state recoding. Box-plots are also generated to visualize the data.

```
perl compare_trees_to_sim_LG.pl
```

**3.9 Simulating under GTR with estimated parameters from the Chang dataset**

**3.9.1** Use the following shell script in the directory 03-SATURATION/04-ESTIMATED_MODEL to reproduce the simulations we performed in Seq-Gen using the Chang topology, amino acid rates of substitution, amino acid frequencies, and gamma rate heterogeneity estimated from the Chang dataset. Simulations were performed with branch length scaling factors 1, 5, 10, 15, and 20.

```
sh run_seqgen_estimated_model.sh
```

**3.9.2** Run the following shell script to create directories for each branch length scaling factor parameter tested and parse the datasets into separate PHYLIP files within their corresponding directories.

```
sh divide_estimate.sh
```

**3.9.3** Create a new directory called 'scripts' and convert the simulated datasets to Dayhoff 6-state recoded datasets using the script chunkify_estimate.pl. chunkify_estimate.pl also generates shell scripts for performing maximum-likelihood analyses in RAxML.

```
mkdir scripts
```

```
perl chunkify_estimate.pl
```

**3.9.5** Execute the shell scripts generated from chunkify_estimate.pl to perform maximum-likelihood analyses on both recoded and non-recoded datasets. We performed maximum-likelihood analyses using the MULTIGAMMA multi-state model with GTR for recoded datasets and the Dayhoff model for non-recoded datasets.

```
ls -1 scripts | grep 'myserver*' | perl -ne 'chomp; print "sh scripts/$_
&\n";' | sh
```

**3.9.6** Run compare_trees_to_sim_estimate.pl to calculate RFD values in TOPD/FMTS for trees produced from recoded and non-recoded datasets and perform t-tests to compare the RFD values between both approaches. This script also generates box-plots to visualize the data.

```
perl compare_trees_to_sim_estimate.pl
```

### 4. Testing Alternative Recoding Strategies on Compositional Heterogeneity

#### 4.1 Test alternative recoding schemes for optimization

**4.1.1** Run the following command on each recoding scheme outlined in Table S1. The script will print a score for each binning strategy specified. Higher scores indicate improved optimization of the Dayhoff matrix.

```
perl score.pl ‘AMINO ACID BINNING SCHEME’
```

#### 4.2 Recode datasets

**4.2.1** Recode the data generated in section 1.3.1 under hypothetical tree 0.002 and inflation parameters 0.1, 0.5, and 0.9 using the best scoring 9-, 12-, 15-, and 18-state binning strategies (Table 2). Perform the following commands in each of the listed directories within 02-COMPOSITIONAL_HETEROGENEITY/02-HETEROGENEOUS_DATA: 07-TREE0002.1, 08-TREE0002.5, and 09-TREE0002.9. Run chunkify_nscheme_recoding.pl to generate shell scripts for maximum-likelihood analyses in RAxML using the MULTIGAMMA multi-state model with GTR.

```
mkdir 9scheme_scripts

mkdir 12scheme_scripts

mkdir 15scheme_scripts

mkdir 18scheme_scripts

perl chunkify_nscheme_recoding.pl 9 9scheme_scripts

perl chunkify_nscheme_recoding.pl 12 12scheme_scripts

perl chunkify_nscheme_recoding.pl 15 15scheme_scripts

perl chunkify_nscheme_recoding.pl 18 18scheme_scripts
```

#### 4.3 Phylogenetic reconstructions

**4.3.1** Execute the following commands in directories 07-TREE0002.1, 08-TREE0002.5, and 09-TREE0002.9 to run the shell scripts generated from chunkify_nscheme_recoding.pl that perform maximum-likelihood analyses on recoded datasets.

```
ls -1 9scheme_scripts | grep 'myserver*'| perl -ne 'chomp; print "sh 9scheme_scripts/$_ &\n";' | sh

ls -1 12scheme_scripts | grep 'myserver*'| perl -ne 'chomp; print "sh 12scheme_scripts/$_ &\n";' | sh

ls -1 15scheme_scripts | grep 'myserver*'| perl -ne 'chomp; print "sh 15scheme_scripts/$_ &\n";' | sh
```

```
ls -1 18scheme_scripts | grep 'myserver*'| perl -ne 'chomp; print "sh 18scheme_scripts/$_ &\n";' | sh
```

**4.4 Score trees**

**4.4.1** Run the following commands in directories 07-TREE0002.1, 08-TREE0002.5, and 09-TREE0002.9 to determine what proportion of trees do not correctly reconstruct a monophyletic group of taxa from clades A and B and a monophyletic group of taxa from clades C and D. The output file identifies incorrectly reconstructed trees and calculates the proportion of incorrect trees produced.

```
perl is_mono.pl DAYHOFF.9 > is_mono_dayhoff9.out

perl is_mono.pl DAYHOFF.12 > is_mono_dayhoff12.out

perl is_mono.pl DAYHOFF.15 > is_mono_dayhoff15.out

perl is_mono.pl DAYHOFF.18 > is_mono_dayhoff18.out
```

**4.5 Generate a bar graph for percentage of incorrect trees**

**4.5.1** Run the script comphet_bargraph_nscheme.R in RStudio to generate a bar graph showing the percentage of incorrect trees reconstructed under each alternative recoding strategy, as well as under Dayhoff 6-state recoding, and non-recoding using the Dayhoff model.

**4.6 Z-tests comparing Dayhoff-18 recoding to non-recoding**

**4.6.1** Run the script ztest_dayhoff_dayhoff18.R in RStudio to perform z-tests on the percentages of incorrect trees reconstructed under the Dayhoff model with non-recoding and the best scoring Dayhoff 18-state recoding strategy.
